# Drawings reveal changes in object memory, but not spatial memory, across time

**DOI:** 10.1101/2024.01.26.577281

**Authors:** Emma Megla, Samuel R. Rosenthal, Wilma A. Bainbridge

## Abstract

Time has an immense influence on our memory. Truncated encoding leads to memory for only the ‘gist’ of an image, and long delays before recall result in generalized memories with few details. Here, we used crowdsourced scoring of hundreds of drawings made from memory after variable encoding (Experiment 1) and retentions of that memory (Experiment 2) to quantify what features of memory content change across time. We found that whereas some features of memory are highly dependent on time, such as the proportion of objects recalled from a scene and false recall for objects not in the original image, spatial memory was highly accurate and relatively independent of time. We also found that we could predict which objects were recalled across time based on the location, meaning, and saliency of the objects. The differential impact of time on object and spatial memory supports a separation of these memory systems.

## Introduction

Our visual memories are the tickets we use to re-experience and re-imagine our lives. Despite the importance and utility of visual memory, it has been well established that the strength of our visual experiences is subject to the whims of time. If we encode an experience for a truncated period, we will largely only have a conceptual or gist-based representation of the experience, with few or no details unique to the experience itself (Ahmad et al., 2017a; M. R. Greene & Oliva, 2009; Oliva, 2005). Similarly, our memories continually fade in their strength since the moment they were encoded, with stretched retention intervals resulting in generalized memories (Harvey, 1986) that are prone to errors (Palmer et al., 2013). However, it is unclear what the actual underlying changes in our memory representations are across time that result in them being better or worse in quality. Does each memory component, such as spatial and object memory, steadily change in unison across time? Or, rather, are different memory components differentially affected by time, with some changing at a faster rate?

Despite overwhelming evidence for improved memory with increased encoding, including for scenes (e.g., Ahmad et al., 2017b; Melcher, 2010) and faces (e.g., McKelvie, 1990; Reynolds & Pezdek, 1992)—indicating important implications for eyewitness testimony (Memon et al., 2003)—few studies have investigated the changes within memory content itself. Studies on visual processing and memory have converged on conceptual information as the first information we extract and encode from a scene (Potter, 2012), with this occurring in the first 100 msec of scene viewing (M. R. Greene & Oliva, 2009). Only after at least 150 msec are any details, such as objects, thought to arrive in memory (Rayner et al., 2009). However, there is very little evidence for how visual details further accumulate into memory, with some evidence for relatively even improvement across time for different features, such as the color, size, and location of objects (Melcher, 2006), but other evidence that spatial memory may be differentially affected by encoding time compared to other features (Tatler et al., 2003). In other words, why our memories improve across encoding following gist extraction is still a large open question in the field.

An expansive literature similarly exists for how our memories worsen as time elapses between encoding and recall (e.g., N. R. Greene & Naveh-Benjamin, 2023; Talamini & Gorree, 2012), including Ebbinghaus’ famous forgetting curve quantifying the rapid rate of verbal forgetting over a century ago (Ebbinghaus, 1885). As our memories age, they are thought to become more generalized (Harvey, 1986; Posner & Keele, 1970; Sweegers & Talamini, 2014; Zeng et al., 2021), leading to the insertion of semantically-related information that was not in the original experience (i.e., *false memories*; Lampinen et al., 2001). This accumulation of incorrect information is accompanied by the loss of details that are tangential to the meaning of an image (e.g., Tompary & Davachi, 2017), providing evidence that the last information to arrive in memory is also the first to leave (Potter, 2012).

This loss of detail over time is not thought to occur in unison—with the color, identity, and location of an object all declining at the same rate—but instead the features of our memories are thought to decline in a predictable (Gold et al., 2005), yet separable fashion. Indeed, studies have even shown that this separation in the forgetting of features occurs early in the course of forgetting (Brady et al., 2013; Gold et al., 2005), such as memory accuracy for the color and the state of an object separable on the scale of minutes after encoding (Brady et al., 2013). However, like studies investigating the effect of encoding time on memory, the relationship of features across the course of forgetting is not well-defined. Most reported dissociations across forgetting are similarly between object and spatial memory, though the nature of this dissociation remains unclear. Whereas there is evidence that spatial memory continues to decline after object memory has stabilized (Talamini & Gorree, 2012), there is also evidence that spatial memory may have an advantage over the course of forgetting, remaining intact even months later (Mandler & Ritchey, 1977). In other words, though there has arguably been more groundwork laid for the course of forgetting than for encoding, a clear picture has yet to be formed for the features of memory driving the quality of our representations across time.

However, drawing has recently been established as a quantifiable method of memory recall that is able to richly capture the content of our memories on the feature level (Bainbridge, 2021). Indeed, drawing can capture the specific objects in memory alongside their associated detail and spatial information, as well as the presence of any false memories. In fact, the rich detail afforded by having people draw what they remember has been able to characterize memory in unique ways beyond other methods, such as the recognition methods classically used in studies on time and memory (Bainbridge et al., 2019; Bainbridge, Pounder, et al., 2021; Hall et al., 2021). Recognition studies often test on images as a whole, meaning that it is difficult to know exactly what aspects of an image are driving performance. Even in recognition studies that probe memory of individual objects or features (like location), novel lures must be used, which requires assumptions on how information is changing in memory when selecting those lures. However, having someone draw what they remember allows for detailed quantification of their memory content without the use of cues or lures. Therefore, drawing-based recall may better pinpoint changes in memory features across time, such as for the accumulating evidence of differential processing of objects and their spatial information (Carlesimo et al., 2001; Reagh & Yassa, 2014; Talamini & Gorree, 2012).

In the current study, we leveraged this detailed window into memory to capture changes in memory across time by having participants draw what they remember from a scene across variable encoding durations (Experiment 1) and delays between encoding and recall (Experiment 2), resulting in hundreds of drawings representative of what is stored in memory across time. To quantify the content of these memory representations, we had roughly 1600 participants score these drawings online across the two experiments. These quantified drawings revealed that some features of memory are highly influenced by both encoding and delay time, with the proportion of objects recalled from a scene and the presence of false objects predicted by time. However, we found that other features of memory, namely spatial memory, were largely unaffected by either encoding or delay time, supporting a dissociation between object and spatial memory. By comparing the identities of objects drawn across time, we also found that whereas the location and meaning of an object influenced when an object would be drawn across encoding, the saliency of an object determined when an object would be recalled across delay.

## Experiment 1

The goal of Experiment 1 was to quantify how encoding time impacts the content and accuracy of our visual memories by manipulating how long participants viewed scene images.

## Methods

### Participants

One hundred and twenty adults originally participated in Experiment 1 on the online crowd-sourcing platform Amazon Mechanical Turk (AMT). As previous work has found meaningful results with 30 participants per drawing condition (Bainbridge et al., 2019), we aimed to recruit over double this sample size per time condition to account for potential exclusions. We excluded participants’ data if at least half of their drawings were unrelated to the task (e.g., a drawing of an animal), of low effort (e.g., scribbles), and / or not created. Based on these criteria, data from 49 participants were excluded. Therefore, the data from 71 participants (25 female, 3 unreported, average age = 36.4, *SD* = 11.3) were used in the analyses. As different methods of producing the drawings could influence the drawing quality (e.g., it is harder to draw using a trackpad than a computer mouse), only participants with a computer mouse were recruited. Participants were consented following the procedures of the University of Chicago Institutional Review Board (IRB19-1395) and were compensated at a rate of $8/hr, with a base of $0.75 for their time.

To promote increased effort in the task, participants were also incentivized with a $0.01 for every object drawn; on average, participants received a $0.11 bonus based on their performance. To further ensure adequate data quality and performance on the task, only AMT workers with at least a 95% approval rate and prior approval of at least 50 tasks were able to participate. Further, to ensure adequate comprehension of the task, only workers with IP address within the United States were able to participate.

Due to technical errors of converting some of the drawings from base64 code into images, 6 of the drawings from the participants were discarded as they cutoff partway through the image. Additionally, the experiment discontinued partway through for two participants, resulting in only five out of six drawings for those participants. This resulted in a total of 418 drawings, with at least 68 drawings from each time condition (*M* = 69.7). To quantify the drawings into data, 1,264 separate participants scored the drawings on AMT across three scoring experiments (see *Scoring Experiments and Analyses*). To participate in the scoring experiments, AMT workers were only recruited if they met the same requirements as the original drawing experiment to ensure adequate data quality.

### Stimuli

Each participant viewed six images of different scenes, with half of the images of indoor scenes (bedroom, kitchen, and living room) and half of outdoor scenes (amusement park, park, and garden). These scene images originate from the SUN database (Xiao et al., 2010) and were selected as they have been shown to have high recall success in previous drawing studies (Bainbridge et al., 2019). Additionally, as studies have found consistency in what images we tend to remember in recognition tasks (i.e., an image’s *memorability*; Bainbridge et al., 2013; Isola et al., 2013), half of the images chosen had low memorability and the other half high memorability—as verified through previous large-scale continuous recognition memorability experiments (Bylinskii et al., 2022; Isola et al., 2011)—although there was no expectation of memorability affecting how well the images were recalled (Bainbridge et al., 2019).

Additionally, all objects within each of the six scenes were annotated with red outlines using the LabelMe Toolbox (Russell et al., 2008) to allow for easy crowdsourced scoring of the objects present in the drawings and to gather ground truth spatial coordinates for the objects in the images. For these annotations, objects were defined as easily nameable, visually distinct objects separable from the rest of the scene. Only whole, separable objects were counted (e.g., a chair leg was not counted as a separate object from a chair, but a curtain was counted as a separate object from a window). Objects less than 50 pixels in diameter were also not counted.

### Procedure

The experiment consisted of six interleaved encoding and drawing recall phases (see **Fig. 1a**). To avoid any differences between encoding the first image and subsequent images, participants were informed that they would be drawing the images from memory before beginning the experiment. The encoding phase began following a 500 msec fixation cross followed by a single image. Each of the images were shown for a different length of time (100 msec, 200 msec, 500 msec, 1 sec, 5 sec, or 10 sec), with participants unaware of how long each image would be presented. Both the order of stimulus presentation and their corresponding encoding durations were randomized across participants. Images were presented at 500 × 500 pixels. The drawing recall phase, in which participants drew what they could remember of the scene, began immediately following encoding. During the drawing recall phase, the drawing canvas matched the size of the image (i.e., 500 × 500 pixels). Participants had large control over their drawings, with access to a variety of mouse colors, an undo tool, and an erasure tool. Participants had unlimited time to create the drawings, but they had to draw for a minimum of 30 seconds before they could continue to the next encoding phase. On average, participants spent 54.2 seconds on each drawing and 33.78 minutes to complete the entire task.

**Figure 1.**
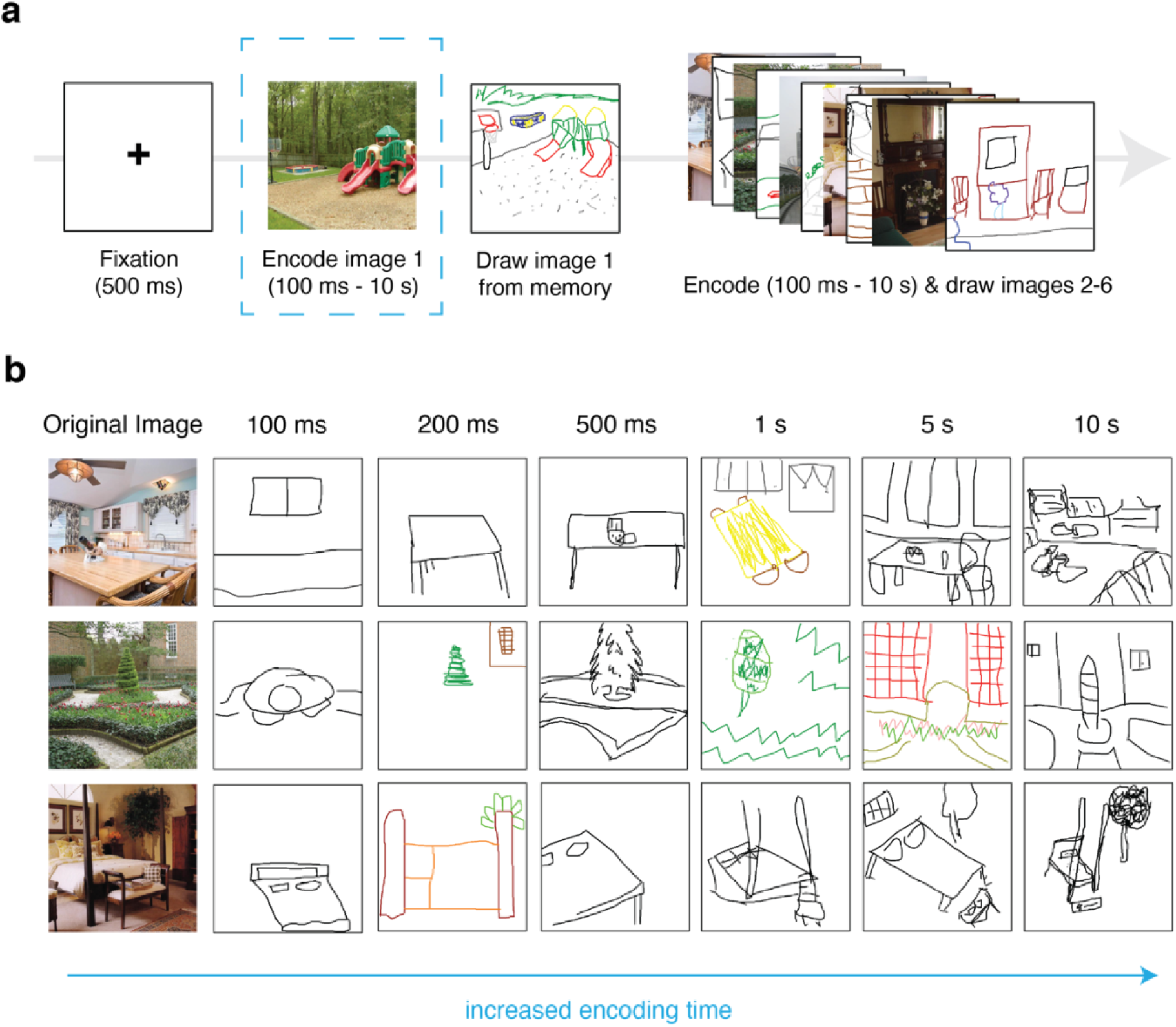
Experiment 1 methods and example drawings. (a) Each trial was preceded by a 500 msec central fixation. After fixation, a single scene image was presented on the screen to encode into memory. To determine the effect of encoding time on what participants can recall from memory, encoding time varied within-participants, with a scene encoded for either 100 msec, 200 msec, 500 msec, 1 sec, 5 sec, or 10 sec. After encoding, participants immediately drew what they could recall from the scene. Each participant completed six trials of this task, encoding each scene for one of the unique encoding durations. (b) Example drawings from the experiment across each of the encoding durations. Although each participant recalled one scene from each of these encoding times, these example drawings are between-participants.

### Online Scoring Experiments

The initial drawing study resulted in drawings representative of what is stored in memory across different encoding times (see **Fig. 1b**). To quantify the content of these drawings, the drawings were run through three online scoring experiments on AMT to determine (1) the objects present in each drawing, (2) the location and size of these objects, and (3) whether the drawings contained any objects not in the original image (i.e., false objects). Participants could complete as many scoring trials as they wanted, and were compensated approximately $6/hr, or $0.01 – $0.03 per trial). A total of 1,263 participants helped score the drawings across the three experiments, which were coded using HTML and JavaScript. These scoring experiments have been widely used to quantify drawings in previous drawing recall studies (e.g., Bainbridge et al., 2019, 2021; Hall et al., 2021). Blank drawings and drawings of the fixation cross were not scored and were counted as having no correctly recalled objects or false objects.

#### Object selection online experiment

We first had AMT workers score which objects were in each drawing. For each drawing, participants were presented with five versions of the original image, each with a different object outlined in red, and were asked to select which objects were present in the drawing. To obtain accurate ratings, five participants rated each object in each drawing. If at least three participants selected the object, it was counted as present in the drawing. As each image contained a different number of objects, we report recall accuracy as the *proportion* of objects recalled from a scene, which is less influenced by scene type. In total, 873 unique AMT workers participated in this experiment.

#### Object location online experiment

For every object deemed present in each drawing, we had AMT workers determine the size and location of the object. Participants were presented with the original image, in which the target object was outlined in red, alongside the drawing. We instructed participants to move and resize a filled ellipse to cover the drawn version of the outlined object in the image. Five participants did this for each object, and we took the median rated spatial coordinates of the ellipse to protect against incorrect ratings. Specifically, we took four spatial coordinates from the ellipse: its central coordinates (both *x* and *y*) determined the location of the object, and its radii (both in *x*- and *y*-direction) determined the size of the object. These spatial coordinates were then compared to the veridical coordinates of the object in the original image by subtracting the true coordinates from the drawn coordinates. Then, we averaged across these error values for every recalled object in each drawing to obtain four measurements of spatial error for each drawing: the *x*-direction location, the *y*-direction location, object width, and object height. These error values were converted from pixels to proportion of the image. In total, 252 unique AMT workers scored the location and size of the objects in the drawings.

#### False objects online experiment

For each drawing, we asked five AMT workers to free respond whether there were any objects drawn that were not in the original image (i.e., *false objects*). We counted a drawn object as a false object if at least two of the five participants responded that a given object was present. In total, 139 AMT workers participated in this experiment. To determine trends in the locations of these false objects, we moved and resized a filled ellipse to cover each false object within every drawing. We then overlaid these ellipses to form heatmaps representing the density of false objects at every pixel.

### Analyses

#### Saliency and meaning maps

The saliency and meaning maps for each scene image were originally created by Bainbridge and colleagues (2019). In brief, to create the saliency maps, each scene was run through a MATLAB toolbox (Harel et al., 2006), using graph-based visual saliency (GBVS). This algorithm determines the most visually dissimilar areas (i.e., the most *salient* areas) of an image, resulting in pixel-level values indicating the dissimilarity of that pixel to the rest of the image. All the pixels within each object of a scene were averaged, which resulted in a single ‘saliency’ value for each object. Meaning maps were created following the conventions of Henderson and Hayes (2017), in which participants rated how “meaningful,” or how informative, an isolated circular patch of a scene was. Therefore, these ratings captured semantic content of the images, rather than just their visual properties. Each scene contained thousands of isolated patches, sampled at both small (3°) and large (7°) sizes, and each participant rated 60 patches. The average meaning scores of these patches were then smoothed, averaged, and combined, resulting in pixel-level ratings indicating the meaning of each pixel within the scene. All the pixels within each object were averaged, resulting in a single ‘meaning’ value for each object.

#### Logistic regression model

An object’s meaning and saliency has been shown to guide what people remember in a scene (Bainbridge et al., 2019). Therefore, we generated a model to determine whether these same features are predictive of whether an object is recalled from a scene across encoding times. The saliency and meaning of each object were determined through the creation of saliency and meaning maps (see *Saliency and meaning maps*). Since the outcome of the model is binary— with an object either drawn or not drawn—we chose a logistic regression model. As we were especially interested in whether salient objects and meaningful objects were more likely to be recalled at a specific point over the course of encoding, we included the interactions of these object features with time along with their main effects.

Since modeling was on the individual object level, we included all objects—irrespective of scene category—in the same model. Therefore, we first included three random effects in the model: a random effect of participant, a random effect of drawing number, and a random effect of scene image (e.g., kitchen, bedroom) We then chose our final model based on the lowest Akaike information criterion (AIC) value—a measure of model fit that takes into account the number of predictors—while still including our main effects of encoding time, object saliency, and object meaning as predictors. As the model with only drawing as a random effect had the lowest AIC value (AIC = 51578, next lowest AIC = 51847), we chose this as our final model. All values were z-scored before being entered into the model.

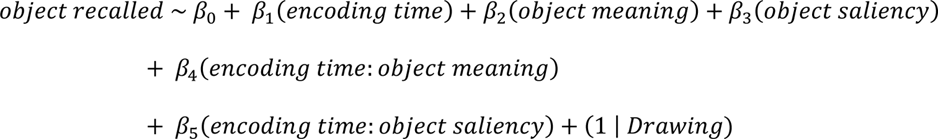

However, because the saliency and meaning of the objects were highly correlated (Spearman’s rank correlation: ρ = 0.84, p <0.001), we ran two additional models to eliminate this multicollinearity. In one, we predicted an object’s meaning from an object’s saliency (*object meaning* ∼ β_0_ + β_1_(*object saliency*)), and in the second, we predicted an object’s saliency from an object’s meaning (*object saliency* ∼ β_0_ + β_1_(*object meaning*)). As the residuals of these models are the variance of each feature that was unexplained by the other, we used these residuals as our saliency and meaning values in our main logistic regression. In other words, any significant effects of saliency and meaning are due to their unique variance, rather than any shared variance.

## Results

### Memories contain more objects over the course of encoding

The longer we encode an image, the more we can “take in” its details, resulting in better memory. For scene images, this may include capturing more objects from the scene into memory. To quantify whether participants were able to recall more objects from a scene after longer encoding, 5 AMT workers (873 workers total) rated whether each object from the original image was present in each drawing.

To determine whether encoding time affects object memory, we ran a mixed-effects linear regression predicting the *proportion of objects recalled from a scene* with *encoding time* (in msec) as a predictor and *participant improvement* (i.e., encoding time | participant) as a random slope (see **Fig. 2a**). We found that longer encoding does indeed lead to more objects in memory with both overall significance of the model (p <0.001, adjusted *R^2^* = 0.34) as well as encoding time as a significant positive predictor of object memory (p <0.001; see *Supplemental Table 1.1* for more detailed statistics for all Experiment 1 models). Indeed, participants could recall a significantly higher proportion of objects at 10 sec (*M* = 0.16, *SD* = 0.12) than 100 msec of encoding (*M* = 0.06, *SD* = 0.07; paired sample *t*-test: *t*(69) = 7.02, *p* <0.001), with participants on average recalling an additional 10.74% of objects from the image (*SD* = 12.7%) over the course of encoding within-participants (see **Fig. 2b** for examples of drawings within participants). When comparing the average proportion of objects participants were able to recall at each time point, we found the largest improvement in object memory when first encoding an image, with the largest increase per millisecond of encoding between 100 msec and 200 msec.

**Figure 2.**
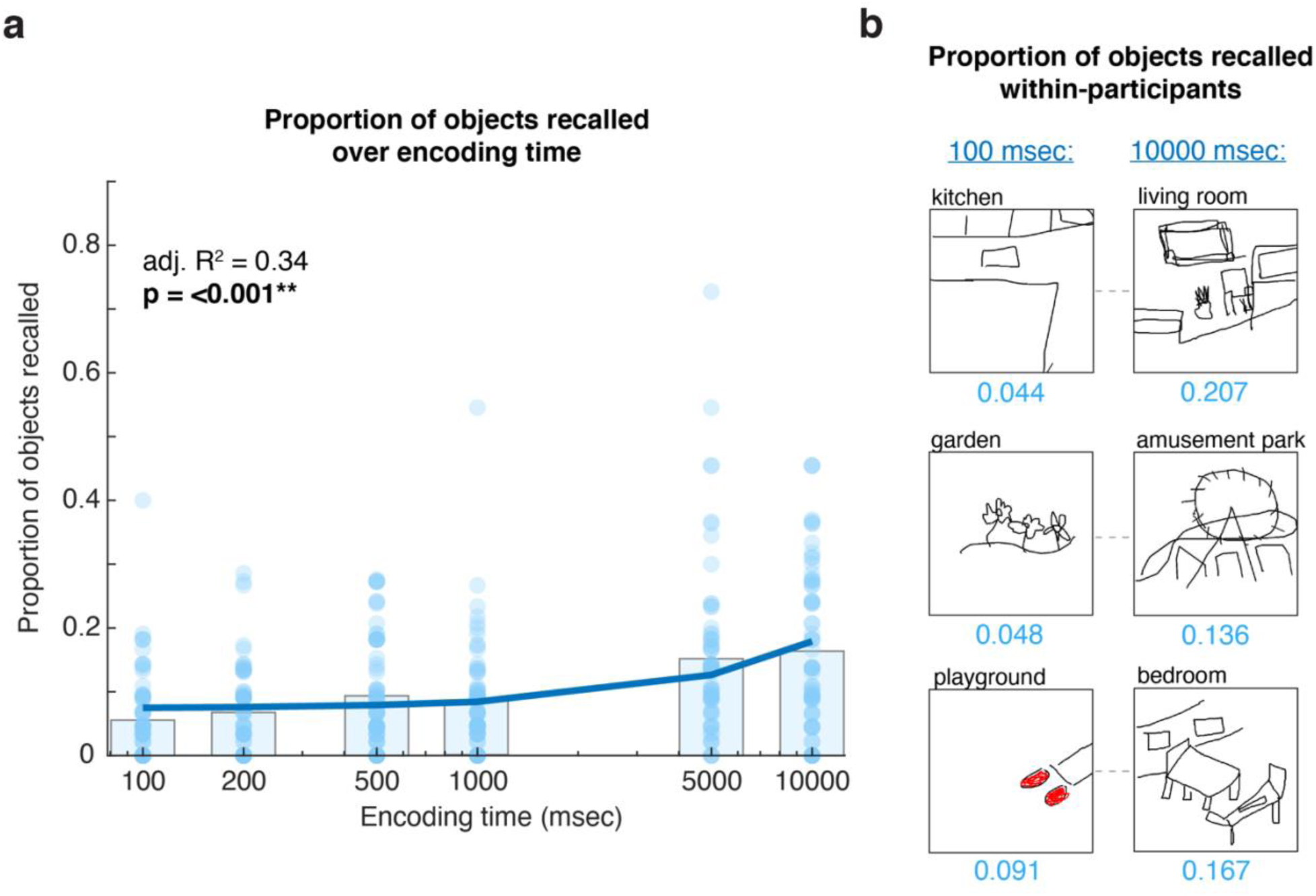
Proportion of objects recalled across encoding time. (a) Encoding time significantly predicted the proportion of objects recalled in a mixed-effects linear regression, with participants recalling significantly more objects with longer encoding. Each light blue dot is the proportion of objects recalled from each drawing, whereas the gray bars are the averages at each time point. A logarithmic scale is used for the x-axis. (b) Example drawings within-participants from the two extreme encoding durations.

On average, participants recalled an additional 0.012% of objects per millisecond (*SD* = 0.099%) across this time interval, with the second highest rate of improvement between 200 msec and 500 msec (*M* = 0.008% per millisecond, *SD* = 0.032%). Therefore, these results suggest that an increase in object memory at least in part contributes to the boost in memory from longer encoding, especially between shorter encoding durations.

### Memories contain different objects depending on how long the scene was encoded

In addition to recalling more objects across encoding, are there also differences in *which* objects are recalled across time? To visualize potential differences in which objects are recalled across encoding, we created heatmaps by subtracting the proportion of drawings containing each object at 10 sec versus 100 msec of encoding for each scene (see **Fig. 3a**). The variability in these heatmaps—with objects ranging from deep blue (i.e., more likely to be recalled after 100 msec of encoding) to deep red (i.e., more likely to be recalled after 10 sec of encoding)—indicates that there do indeed appear to be trends in which objects are recalled across encoding.

**Figure 3.**
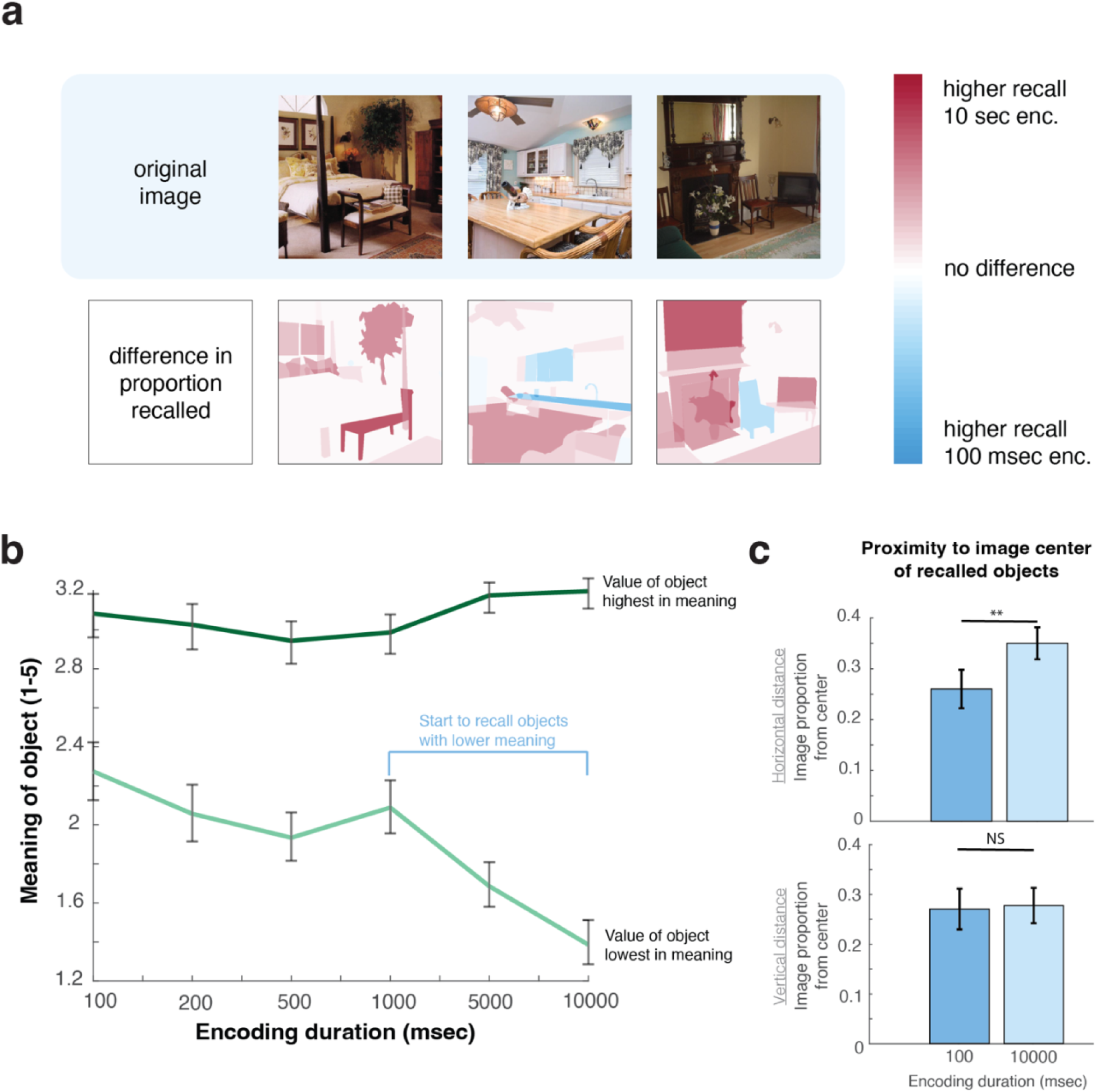
The differences in objects recalled across encoding. (a) Object heatmaps created by subtracting the proportion of participants that recalled an object at 10 sec of encoding minus 100 msec of encoding. The variability within these heatmaps indicate that there are likely object properties that predict their encoding and recall across encoding time. (b) The average highest and lowest meaning value of an object in drawings across time. The dip in the minimum meaning value in longer encoding periods reflects the finding of objects lower in meaning being encoded after more meaningful objects (c) Bar graphs comparing the proximity to image center of recalled objects at 100 msec versus 10 sec. Participants recalled objects significantly closer to the horizontal center, but not vertical center, of the images at 100 msec compared to 10 sec within-participants. Error bars in all figures are standard error of the mean.

To pinpoint which object properties predict their recall across encoding, we analyzed the effect of object meaning and saliency (see **Fig. 3b**) as well as the location of the objects (see **Fig. 3c**) across time. First, we ran a logistic regression model with the main effects of *encoding time*, *object meaning*, and *object saliency.* The interactions of *encoding time and meaning* and *encoding time and saliency* were also included as predictors in the model in addition to the random effect of the *individual drawing* (see *Methods* for further details about the model).

Overall, we found that this regression model could significantly predict whether an object was drawn (p <0.001, adjusted *R*^2^ = 0.05; see *Supplemental Table 1.2* for additional model statistics). Additionally, as expected, we found that longer encoding increased the likelihood of an object being drawn (p <0.001), as well as higher saliency (p <0.001) and higher meaning (p <0.001) of the object. However, we only found that an object’s meaning predicted when an object would be drawn across encoding durations (p = 0.008), with an object high in meaning more likely to be recalled after short encoding durations. The saliency of an object had no influence on when an object was drawn across time (p = 0.781). In other words, it seems that an object’s meaning, rather than its saliency, may have a larger sway in what we first choose to encode during scene perception. To verify this visually, we also determined the objects highest and lowest in meaning within each drawing and visualized how these values changed across time (see **Fig. 3b**). We found that although the maximum meaning value remained relatively stable across encoding, there was a large decrease in the minimum meaning value with longer encoding. In other words, unlike meaningful objects, objects low in meaning were not encoded until further in the course of perceptual processing.

Additionally, previous work has indicated a central bias during encoding (Tatler, 2007), meaning that objects in the center of scenes may be the first objects to enter into memory. To test whether this bias influenced which objects were recalled under short encoding times, we calculated how far each drawn object was from the image center for every scene. Specifically, we subtracted the coordinates of the center of the image from the center of each object. We did this calculation separately for the *x*- and *y*-coordinates, then took the absolute values. Then, we averaged across these values for every object in each drawing, resulting in how central the recalled objects were in each drawing in both the *x*- and *y*-directions. To determine whether more centrally located objects were recalled with shorter periods of encoding, we compared the two extreme encoding conditions: 100 msec and 10 sec. From this, we found evidence of a central bias during encoding—objects were recalled significantly closer to the horizontal center of the image (i.e., the *x*-direction) at 100 msec of encoding (*M* = 0.26, *SD* = 0.17) than at 10 sec of encoding (*M* = 0.35, *SD* = 0.14) within-participants (t(34) = −2.89, p = 0.007). However, participants were no less likely to recall objects close to the vertical center (i.e., the *y*-direction) of the image at 100 msec (*M* = 0.27, *SD* = 0.18) than 10 sec of encoding (*M* = 0.28, *SD* = 0.16; t(34) = −0.19, p = 0.85). In other words, we found that objects closer to the horizontal center of the images, but not the vertical center, were prioritized during encoding. These results suggest that specific object properties—such as meaning and location—can predict whether an object will be recalled across encoding.

### Memories become more accurate across encoding

Our memories can be characterized not only by how detailed they are (i.e., the amount of correct information), but also by how *accurate* they are (i.e., the amount of false information). To determine the presence of false memories—or in this case, false objects in drawings—we had 5 AMT workers (N = 139 workers total) view each drawing and free respond whether there were any objects drawn that were not in the original image. Overall, we found that very few drawings contained any false objects (32/418 or 7.65%; see **Fig. 4a** for examples), and that most drawings that contained any only had one (27/32 or 84.38%). To determine whether encoding time predicts the number of false objects in memory, we ran a mixed-effects linear regression predicting the *number of false objects* in a drawing with *encoding time* as a predictor and *participant improvement* as a random slope. We found that increased encoding time led to fewer false objects in memory (see **Fig. 4b**), with significance of the overall model (p = 0.027, adjusted *R*^2^ = 0.13) as well as encoding time as a significant predictor (p = 0.027). In other words, our memories not only become more detailed over the course of encoding, but also more accurate, with fewer false objects.

**Figure 4.**
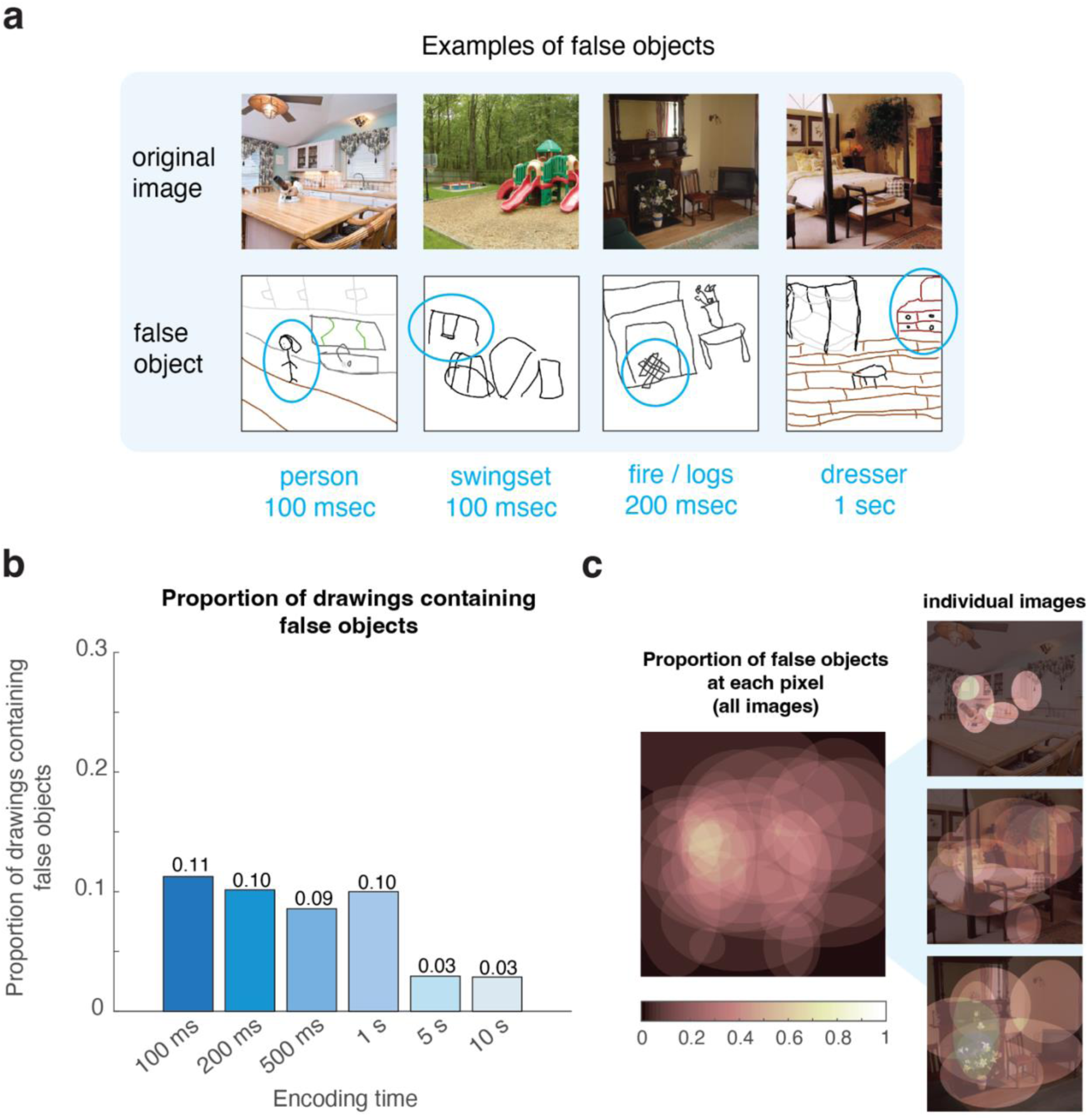
False objects in drawings across variable encoding times. (a) Examples of drawings containing false objects. Participants tended to either falsely recall people or objects related to the scene recalled, such as a dresser in the bedroom. (b) A bar graph comparing the proportion of drawings containing at least one false object recalled across encoding. As confirmed with a separate mixed-effects linear regression, participants significantly recalled fewer false objects as they encoded a scene for longer. (c) Heatmaps overlaying the location of each false object across every drawing, averaged across all images (left) as well as within the drawings for a single image (right). Visual inspection reveals that false objects tended to be recalled close to the center of the drawing, which may reflect the location of objects in the images.

To determine the properties of these false objects, we investigated trends in their identities and their locations within the drawings. Qualitatively, we found trends in the identities of these objects; false objects were often people (6 drawings) or objects semantically related to the scene, such as a dresser in the bedroom (4 drawings) or fire / logs in the fireplace (2 drawings). Additionally, when we overlaid ellipses covering the size and location of all false objects to create a heatmap (see **Fig. 4c**), we surprisingly found that these false objects tend to be centrally located within the drawings. Overlaying these false object ellipses over their original image reveals that these objects tend to have similar placement to those in the true images; in other words, participants may have recalled the location of true objects, but misremembered their identities. Alternatively, the presence of two central areas of high-density (Fig. 4c)—one left of center and one right of center—could potentially reflect that participants stylistically chose to draw false objects along the “rule of thirds” in their drawings. This rule is a well-known and commonly taught strategy for where to place objects in artwork to make it the most visually pleasing and interesting (e.g., Peterson, 2015). Therefore, it is possible that participants added false objects in their drawings for a similar effect.

Overall, the higher presence of false objects after minimal encoding in combination with these false objects tending to be semantically related to their scene are in line with participants first capturing the “gist” or conceptual information of an image during encoding. Therefore, under limited encoding time, participants are only able to capture this high-level information, resulting in the false recall of objects related to the scene.

### Spatial memory is precise at just 100 msec of encoding

The accuracy of memory can be defined not only by *what* is in a memory, but *where* that information is located. Is spatial memory, like object memory, influenced by the length of encoding? To determine the spatial accuracy of memory, 5 AMT workers (252 workers total) moved and resized an ellipse to cover each recalled object in every scene. By comparing the spatial coordinates of these ellipses to the coordinates of the objects in the original image, we were able to gather four measurements of spatial accuracy for each drawing: the *x*-direction location, the *y*-direction location, object width, and object height.

Overall, participants were incredibly accurate for where they recalled objects (see **Fig. 5**). Across encoding conditions, participants on average displaced objects by 9.98% of the image in the *x*-direction (*SD* = 8.04%) and 11.19% (*SD* = 7.30) of the image in the *y*-direction. These error values are far better than if the objects had been randomly placed, with chance calculated to be around 30% (30.9% in *x*-direction, 30.3% in *y*-direction) in previous drawing studies (Bainbridge et al., 2019). Interestingly, we found that encoding time had little effect on the accuracy of location memory. In fact, a mixed-effects linear regression with *participant improvement* as a random slope revealed that *encoding time* did not significantly predict *location memory in the x-direction* (overall model: p = 0.980, adjusted *R*^2^ = 0.02; predictor: p = 0.980). However, when we used these same predictors to predict location memory in the *y*-direction, we did find that *encoding time* negatively predicted the *displacement of objects in the y-direction* (overall model: p = 0.035, adjusted *R*^2^ = 0.04; predictor: p = 0.035), although the improvement across encoding within-participants was minimal (*M* = 1.24%, *SD* = 8.52%).

**Figure 5.**
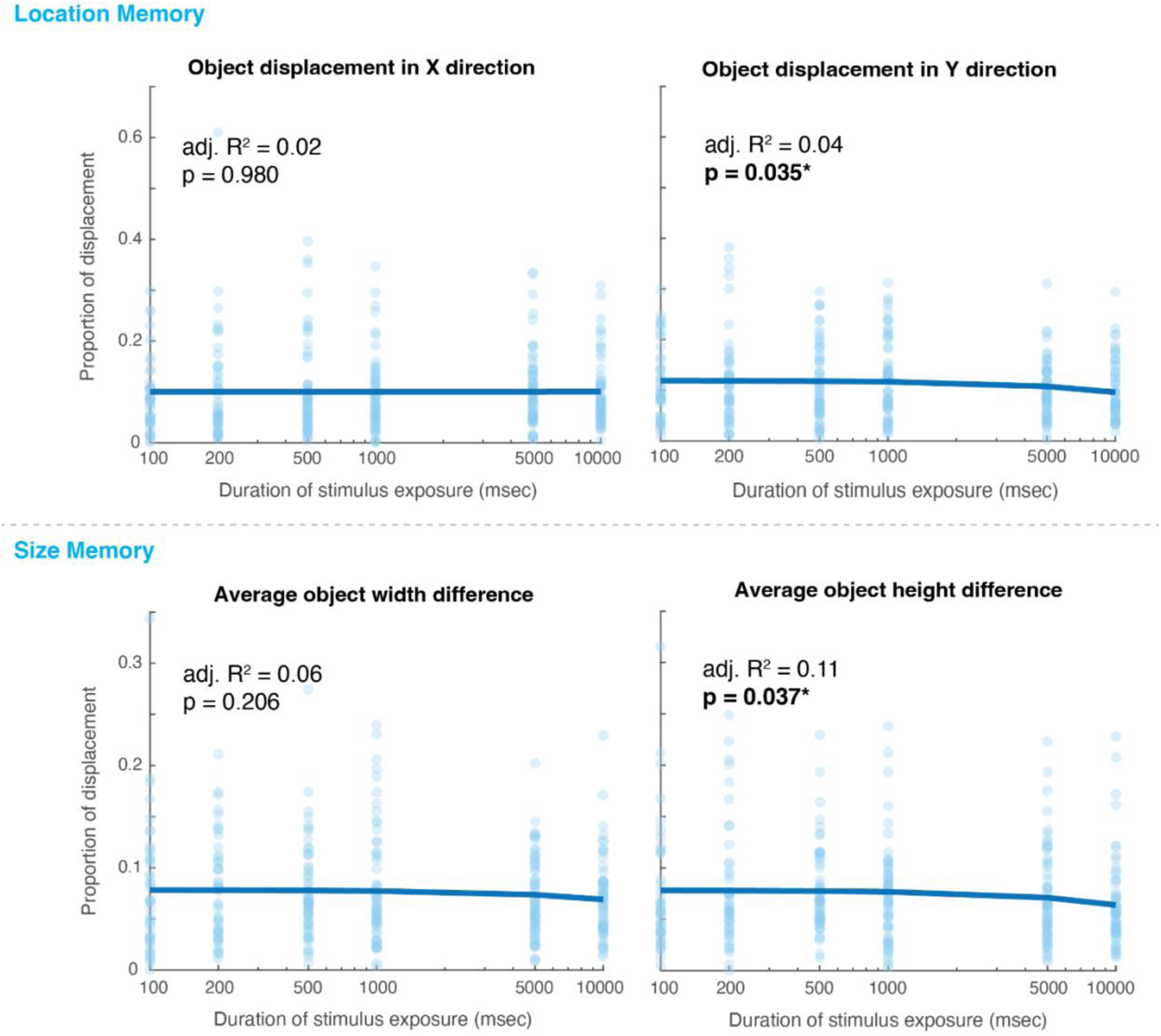
Mixed-effects linear regressions predicting spatial accuracy from encoding time. (top panel) For location memory, encoding time was only significantly predictive of location accuracy in the y-direction, but not the x-direction, meaning that increased encoding time led to improved location memory in the y-direction. (bottom panel) Similarly for size memory, encoding time was only significantly predictive of height memory, not width memory, meaning that increased encoding led to improved memory for how tall an object was.

Similar to location memory, we found that participants were surprisingly accurate in their memory for the size of objects. Across encoding conditions, participants were on average 7.54% (*SD* = 4.92%) of the image off for the width of objects and 7.30% (*SD* = 4.93%) of the image off for the height of objects. Interestingly, we found the same differential effect on the *x*- vs. *y*- directions in size memory, with *encoding time* not predictive of *width memory* (i.e., memory in the *x*-direction) in a mixed-effects linear regression with *participant improvement* as a random slope (overall model: p = 0.206, adjusted *R*^2^ = 0.06; predictor: p = 0.206), but negatively predictive of height memory (i.e., memory in the *y*-direction), meaning that height error decreased with longer encoding, in a separate mixed-effects model with the same predictors (overall model: p = 0.037, adjusted *R*^2^ = 0.11; predictor: p = 0.037). Again, although encoding time was significantly predictive of height memory, the improvement across encoding was minimal within-participants (*M* = 1.35%, *SD* = 7.08%). These results suggest that our spatial memory is overall incredibly accurate, and that it is only minimally—if at all—dependent on how long the image was encoded. These results also provide interesting evidence for a dissociation between our memory for *x*- vs. *y*-coordinates, with only our memory in the *y*- direction minimally improving across encoding time.

Overall, we found that encoding time had differential effects on unique memory components. Whereas participants recalled a higher proportion of objects—and fewer false objects—with longer encoding, memory for the location and size of objects was relatively unaffected by time, and was in fact incredibly accurate at just 100 msec of encoding. Additionally, our results indicate that specific object properties are predictive of what is recalled from memory across encoding, with longer encoding resulting in less meaningful and more peripheral objects drawn from memory.

## Experiment 2

The goal of Experiment 2 was to quantify how the length of the retention period between encoding and recall impacts the content and accuracy of our visual memories. We did this by manipulating how much time elapsed (referred to here as “delay time”) between study and recall of a scene image.

## Methods

### Participants

1,609 adults originally participated in Part 1 (encoding phase) of the experiment on Prolific, with 942 participants (57.5%) returning for Part 2 (drawing recall phase). Participants were continually recruited until we achieved a similar number of drawings as Experiment 1. To screen for potential cheating, we recorded key presses during image encoding and only invited participants back for Part 2 if no keys were pressed. Of the participants that returned, we excluded the data from 65 participants due to not answering the survey questions, not creating a drawing, and / or creating a drawing that was clearly low effort (e.g., scribbles) or off task (e.g., an animal drawing). We also excluded 426 participants as they did not return within the designated time frame (+/- 20% of the time condition; *see Procedure*). An interesting sample of 19 participants (11 female, average age = 33.1, *SD* = 11.4) returned after a retention interval longer than any of our predetermined conditions (returned 1-2 weeks after encoding), and were included in a small portion of analyses. In total, data from 451 participants (215 female, 16 unreported, average age = 32.0, *SD* = 9.6) were used in all analyses for a total of 451 drawings, roughly matching the number of drawings from Experiment 1, with at least 74 drawings in each time condition (*M* = 75.2, *SD* = 1.6). Participants were consented following the procedures of the University of Chicago IRB (IRB19-1395), were compensated a rate of $8.24/hr between the two parts, with a compensation of $0.14 for completion of Part 1 and $1.50 for Part 2. Participants were required to have a computer mouse to participate.

To ensure adequate data quality, only Prolific users with at least a 90% approval rate and 50 previous submissions were recruited. Additionally, we only recruited participants from the United States who were fluent in English. Participants also had to pass two attention checks at the end of the experiment, which consisted of correctly providing the term for a baby dog (i.e., puppy) and the month Halloween takes place (i.e., October). To further protect against cheating, participants also had to respond that they did not cheat during the experiment for their data to be included.

To score the 451 drawings from the main drawing experiment, 1,981 participants on AMT scored the drawings across three scoring experiments. In order to score the drawings, the AMT workers had to meet the same criteria from the scoring experiments of Experiment 1.

### Stimuli

The stimuli were the same as in Experiment 1, but each participant only viewed one of the scene images in Experiment 2.

### Procedure

This experiment consisted of one encoding phase and one drawing recall phase (see **Fig. 6a**). In the encoding phase, participants encoded one of the randomly chosen six scene images for 10 seconds. Participants only encoded one image in this experiment to avoid interference in memory between images. After encoding the image, participants were asked to select which times they would be available to participate in the second phase of the experiment, with six options available (immediately, 5 minutes later, 1 hour later, 1 day later, 2 days later, or 1 week later).

**Figure 6.**
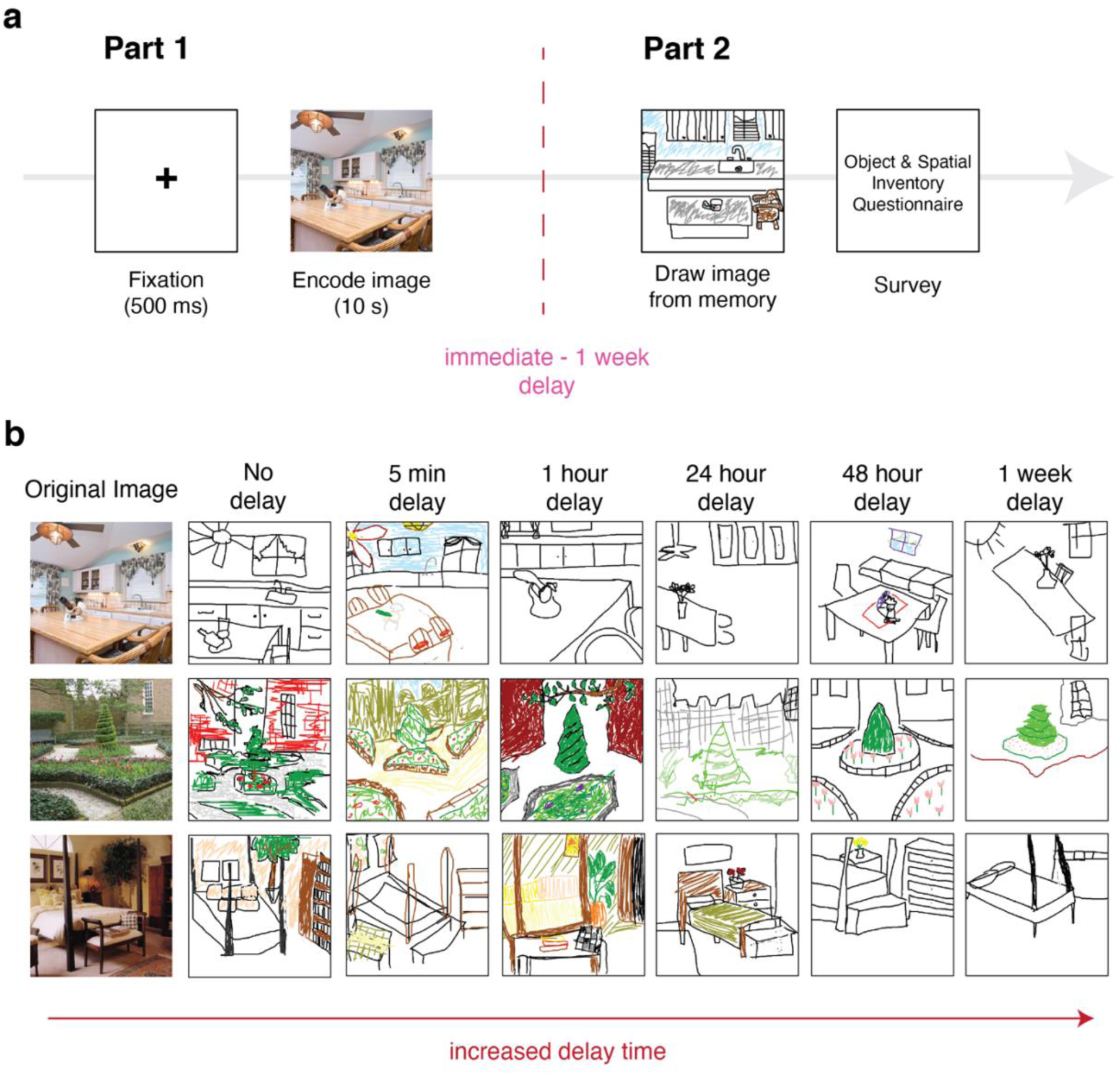
Experiment 2 methods and example drawings. (a) The experiment started with a 500 msec central fixation followed by the presentation of a single scene image to encode for 10 sec. Participants encoded only one of six scene images. Following encoding, participants returned after a delay of either no delay, 5-minutes, 1-hour, 1 day, 2 days, or 1 week to complete Part 2 of the experiment. In the second part of the experiment, participants drew what they could recall from the scene, and then completed the Object & Spatial Inventory Questionnaire (OSIQ). (b) Examples drawings from the experiment across the delay conditions, all drawn by different participants.

After completing the encoding phase, the experimenter assigned the participant one of the selected time frames and messaged the participant with an expiring password and the correct time frame to return within. For each time condition, the time frame to return within was +/- 20% of the time condition. For example, participants in the 5-minute condition could return within 3-6 minutes after encoding.

After returning, the second part of the experiment consisted of a drawing recall phase. In the drawing recall phase, participants were asked to draw what they remembered from the scene image presented in the encoding phase. The drawing canvas, drawing tools, and minimum time requirement were the same as Experiment 1. Following drawing recall, participants were asked to complete the Object and Spatial Inventory Questionnaire (OSIQ; Blajenkova et al., 2006)—a survey that assesses both the vividness of object (i.e., Object score) and spatial mental imagery ability (i.e., Spatial score)—to determine whether these mental imagery abilities influence their respective memory measures. Although it is possible that mental imagery could also influence memory recall across encoding time, we unfortunately did not think of this link during Experiment 1. On average, Part 1 of the experiment took about 1.58 minutes to complete and Part 2 took about 8.05 minutes to complete.

### Scoring experiments

The initial drawing study resulted in drawings representative of what is stored in memory across retention intervals (see **Fig. 6b**). The same scoring experiments from Experiment 1 were used to quantify the content of these drawings. A total of 1,988 AMT workers scored the drawings, with 847 participants scoring the objects recalled, 831 participants scoring the size and location of these recalled objects, and 310 participants scoring the presence and identity of false objects.

### Analyses

We used the same image-based metric maps from Experiment 1, in which a meaning and saliency value was calculated for every object in each image.

#### Logistic regression model

We ran the same logistic regression model as Experiment 1, with delay time in place of encoding time, to determine the effect of saliency and meaning on which objects are recalled across delays in recall.

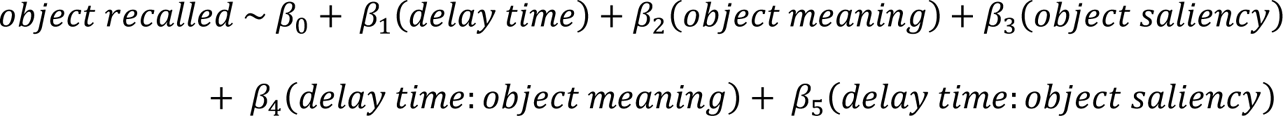

However, because the AIC of the model was lower with no random effect of drawing / participant or scene type (AIC without random effect = 47373, lowest AIC with random effect = 47771), we did not include any random effects in this model.

#### Other statistical methods

When conducting independent samples *t*-tests between delay time conditions, we assumed unequal variance between groups with corrected degrees of freedom.

## Results

### Memories contain fewer objects across longer delays in recall

It is generally thought that the longer a memory is stored, the worse that memory becomes. This may be due to the loss of details in memory, such as fewer objects in our memories for scenes. To test whether participants recalled fewer objects from a scene after longer delays before recall, 5 AMT workers (847 workers total) rated whether each object from the original image was present in each drawing. Using a linear regression, we found that *delay time* (in hours) negatively predicted the *proportion of objects recalled* from a scene (see **Fig. 7a**; F(449) = 29.9, p <0.001, adjusted *R^2^* = 0.060), with fewer objects recalled after longer delays (see *Supplemental Table 1.3* for more detailed statistics for all Experiment 2 models). Indeed, participants who recalled a scene after one week recalled significantly fewer objects (*M* = 0.17, *SD* = 0.17) than those who recalled a scene immediately after encoding (*M* = 0.28, *SD* = 0.17; independent samples *t*-test: t(151.99) = −3.97, p <0.001), losing on average an additional 10.9% of objects over this delay between-participants. As immediate recall could in part be pulling from short-term working memory, rather than the long-term memory store of the one-week condition, we also compared the one-week delay group to the 5-minute delay group. We replicated the same result within the long-term memory conditions, with worse object memory for the one week (*M* = 0.17, *SD* = 0.17) than the 5-minute delay group (*M* = 0.28, *SD* = 0.14; independent samples *t*-test: t(147.16) = 4.204, p <0.001). On average, participants lost an additional 10.7% of objects over the course of this delay between participants. The largest rate of decline occurred between no delay and five minutes of delay, with an average of 0.037% of objects lost per minute in this time frame.

**Figure 7.**
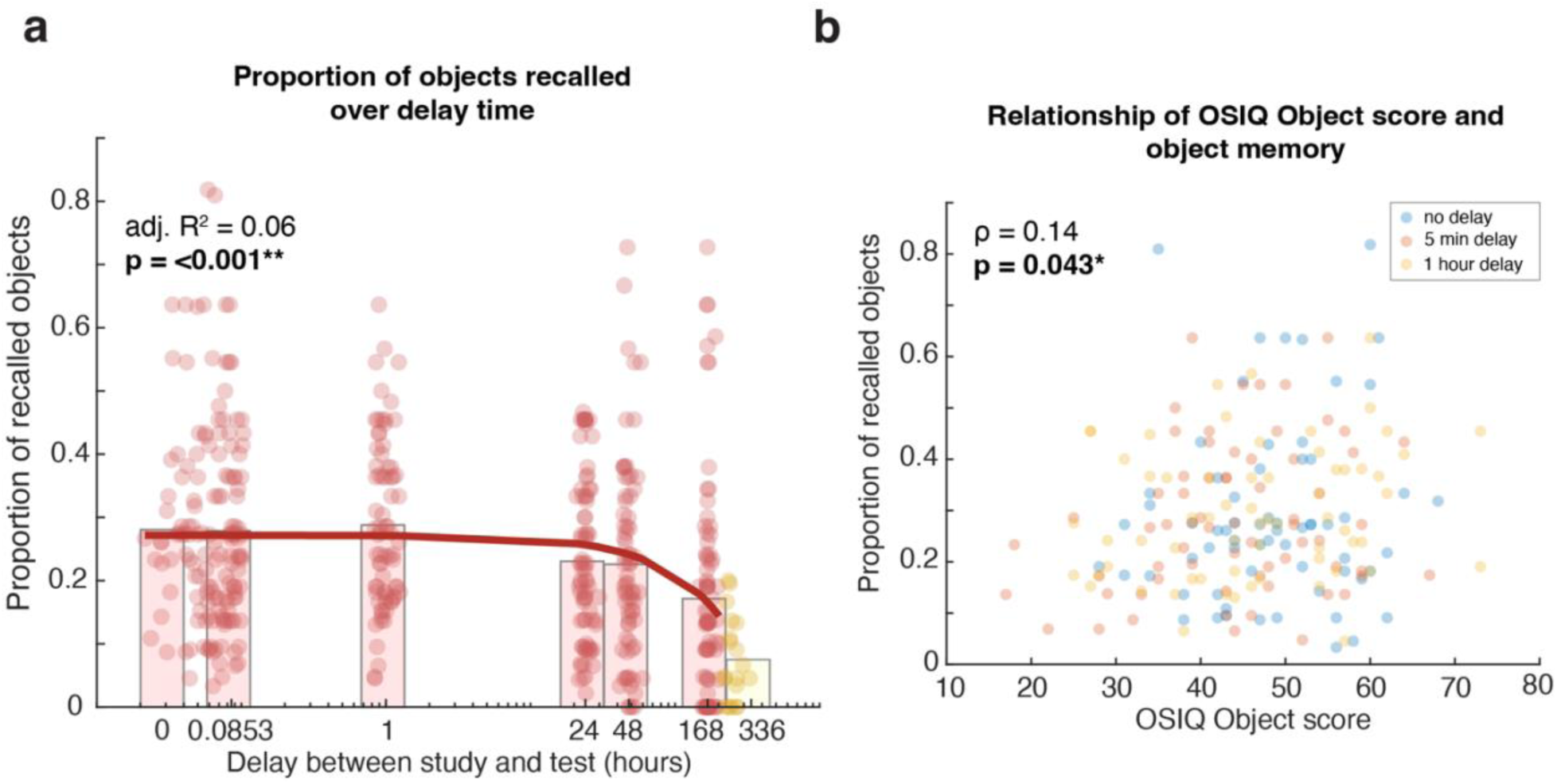
Influence of delay time and OSIQ-Object score on the proportion of objects recalled from a scene. (a) Delay time significantly predicted the proportion of objects recalled in a linear regression, with participants recalling significantly fewer objects after longer delays. Each red dot represents a single scene, whereas the red bars represent the average at each delay time. The yellow dots and bar are from a group of participants that returned between 1-2 weeks after encoding, but these were not included in the actual regression analysis. (b) Correlation between participants’ OSIQ-Object score and the proportion of objects recalled from a scene. To minimize the influence of time on object recall, only participants with the shortest delay times (no delay – 1 hour delay) were included in this correlation. There was a significant correlation between object mental imagery ability (i.e., Object score) and the proportion of objects recalled from a scene, with stronger imagery ability leading to a higher proportion of objects recalled.

However, despite the forgetting that occurred over the 1-week period, participants had surprisingly good memory for their encoded scene. On average, participants were still able to recall on average 17.2% (*SD* = 17.2%) of objects from a scene even one week after encoding.

As the vividness of mental imagery has been shown to impact memory recall (e.g., *aphantasia*; Bainbridge, Pounder, et al., 2021), we also had participants respond to the Object and Spatial Inventory Questionnaire (OSIQ; Blajenkova et al., 2006) at the end of the experiment. Therefore, it is possible that object and spatial imagery may explain a portion of variance in their respective, separable memory measures in the current study. To determine whether object imagery ability influences the proportion of objects recalled, we added participants’ *Object score* as an additional predictor to the previous linear regression predicting the *proportion of objects recalled* from *delay time*. As the AIC of the model was lower without the interaction term in the model (AIC = 1060 with interaction, AIC = −1058 without interaction), the interaction term was not included. However, the model statistics are comparable across both models (see *Supplemental Table 1.4* for model comparison and more detailed statistics). Additionally, only participants that completed the entirety of the OSIQ were included in the model and subsequent OSIQ analyses (385/451 or 85.4% of participants). Overall, the model was significantly predictive of object memory (F(382) = 20.3, p <0.001, adjusted *R^2^* = 0.09), with delay time still a significant negative predictor (p <0.001). Interestingly, we also found that *Object score* did additionally positively predict *object memory* (p = 0.034), explaining additional variance beyond delay time. This result was confirmed in a separate analysis correlating Object score and object memory (see **Fig. 7b**). In other words, both longer delays in recall as well as lower object imagery ability significantly reduce the proportion of objects recalled from a scene.

### Visual saliency predicts which objects are recalled across delays between encoding and test

To visualize potential differences in which objects are recalled across encoding, we created heatmaps subtracting the proportion of drawings containing each object at immediate recall versus 1-week delayed recall for each scene (see **Fig. 8a**). Visual inspection of these heatmaps suggests that there may be object properties that predict which objects are recalled across delay time.

**Figure 8.**
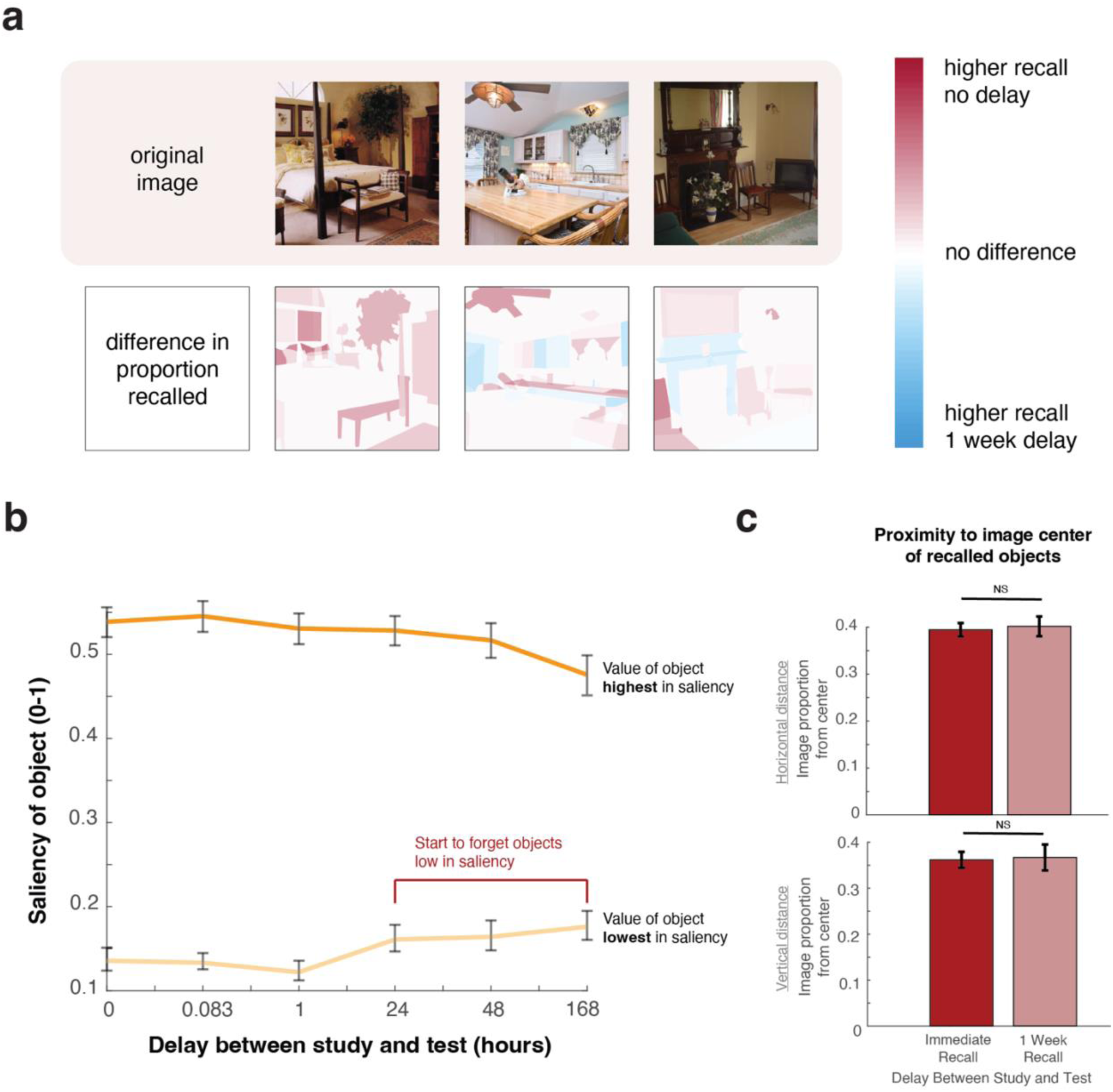
The differences in objects recalled across delays between study and test. (a) Object heatmaps created by subtracting the proportion of participants that recalled an object immediately after encoding minus 1-week after encoding. The variability within these heatmaps indicate that there may be object properties that predict their encoding and recall across encoding time. (b) The average highest and lowest saliency value of an object in drawings across time. The rise in the minimum saliency value in a drawing in the longer delay periods reflects the finding of objects low in saliency being the first objects forgotten from an image. (c) Bar graphs comparing the proximity to image center of recalled objects at immediate vs. 1-week delayed recall. Participants did not recall objects any closer to the horizontal or vertical centers of an image in these extreme delay conditions. Error bars in all figures are standard error of the mean.

To determine the object properties that influence when an object is able to be successfully recalled across delay time, we analyzed the effect of the meaning and saliency of an object (see **Fig. 8a**), as well as the location of an object within a scene (see **Fig. 8c**). First, to test whether the meaning and saliency of an object predicts whether it will be forgotten over time, we ran a logistic regression model with the main effects of *delay time*, *object meaning*, and *object saliency* as predictors for whether an object is drawn. Critically, we also included the interactions of *delay time and meaning*, and *delay time and saliency* as predictors (see *Methods* for further details about the model).

Overall, we were able to significantly predict whether an object was drawn using the combination of these predictors (F(10474) = 40.07, p <0.001, adjusted *R*^2^ = 0.02; see *Supplemental Table 1.5* for more detailed statistics). As expected, we found that increased time between encoding and recall led to a lower likelihood of recalling the object (p <0.001), and that higher saliency and meaning of the objects increased the likelihood of recalling the object (saliency: p <0.001; meaning: p = 0.016. Interestingly, we found that objects lower in visual saliency were more likely to be forgotten after longer delays (p = 0.023), but that the meaning of an object had no effect on whether it would be recalled across time (p = 0.484). These results suggest that whereas meaningful objects are retained in memory across a 1-week delay, objects low in saliency are forgotten. To verify this, we determined the objects highest and lowest in saliency within each drawing and visualized how these values changed across time (see **Fig. 8b**). We similarly found that the minimum saliency value increased across longer time scales, again suggesting that the while high saliency objects stayed in memory, the objects with the lowest saliency were forgotten over longer delay times. Notably, there is also a small, steady dip in the value of the most salient object within a drawing, suggesting that objects across the spectrum of saliency levels may additionally be minimally forgotten across time, though not to the extent of objects lower in saliency.

Additionally, it is possible that the location of an object could influence how well an object is recalled across delays in recall, with the largest distinction in location being how central an object is within an image. To test whether a central bias contributes to what is recalled across delay time, we averaged across the distance from center for all objects recalled within a drawing. However, we found no evidence of this distinction in object location contributing to what is recalled across delay time, with participants recalling objects just as close to the horizontal center by image proportion at immediate recall (*M* = 0.40, *SD* = 0.12) as 1-week recall (*M* = 0.403, *SD* = 0.16) between-participants (independent samples *t*-test: t(108.3) = −0.285, p = 0.776). The same was true for recalling objects close to the vertical center by image proportion (immediate recall: *M* = 0.34, *SD* = 0.13, 1-week recall: *M* = 0.35, *SD* = 0.15; independent samples *t*-test: t(117.6) = −0.358, p = 0.721). Overall, these results suggest that the saliency of an object, rather than its location or meaning, is the most predictive of whether an object will be forgotten across time, with objects low in saliency the most prone to forgetting.

### Memories become less accurate across time

As our memories could also become less accurate (i.e., with more false objects) with delays between encoding and recall in addition to less detailed, we had 5 AMT workers (310 total workers) identify whether there were any false objects in each drawing. Overall, we found that 65/451 (14.4%) drawings contained at least one false object (see **Fig. 9a** for examples), with most of these drawings only containing one false object (49/65 or 75.4%). Participants were more likely to recall false objects after longer delays (see **Fig. 9b**), with *delay time* a significant positive predictor (p <0.001) of the *number of false objects* in a drawing (F(449) = 11.4, p <0.001, adjusted *R*^2^ = 0.02). In line with increased recall of false objects over time, about a fifth of participants that returned around two-weeks after encoding drew at least one false object (4/19 or 21.1%), with 75.0% of these drawings containing more than one (M = 2.25 false objects). In other words, memories not only become less detailed over delays in recall, but also less accurate.

**Figure 9.**
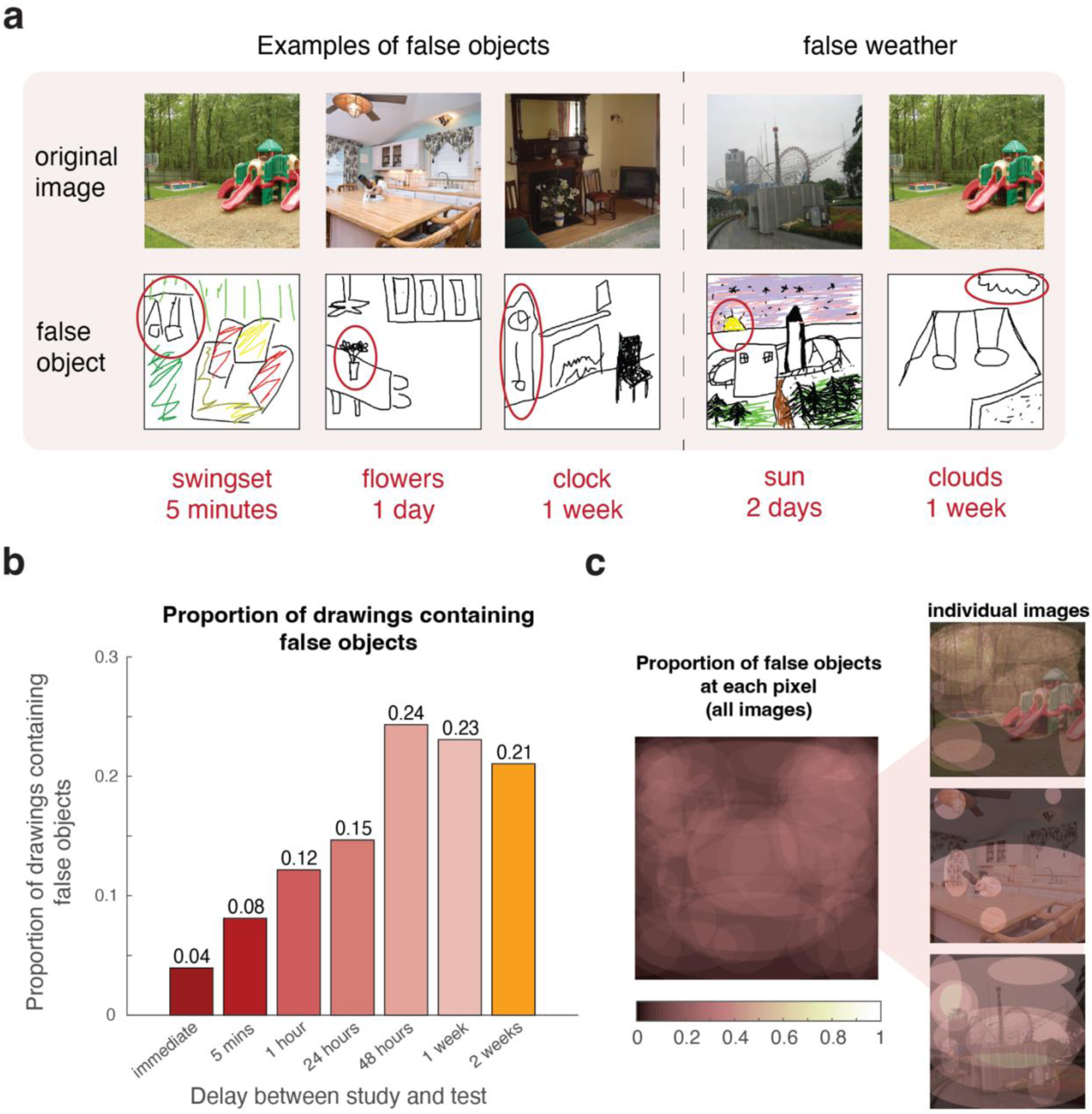
False objects in drawings across variable delays before recall. (a) Examples of drawings containing false objects. Participants tended to falsely recall objects related to the scene recalled, such a swingset in the playground, or false sceneries, such as a sun in the sky. (b) A bar graph comparing the proportion of drawings containing any false objects across delay times. As confirmed with a separate linear regression, participants significantly recalled more false objects as more time elapsed before recalling the encoded scene. (c) Heatmaps overlaying the location of each false object across every drawing, regardless of image (left) as well as within the drawings for a single image (right). Visual inspection reveals that though false objects occasionally overlapped with the location of true objects in the recalled image, overall there were minimal observable trends for the location of false objects.

**Figure 10.**
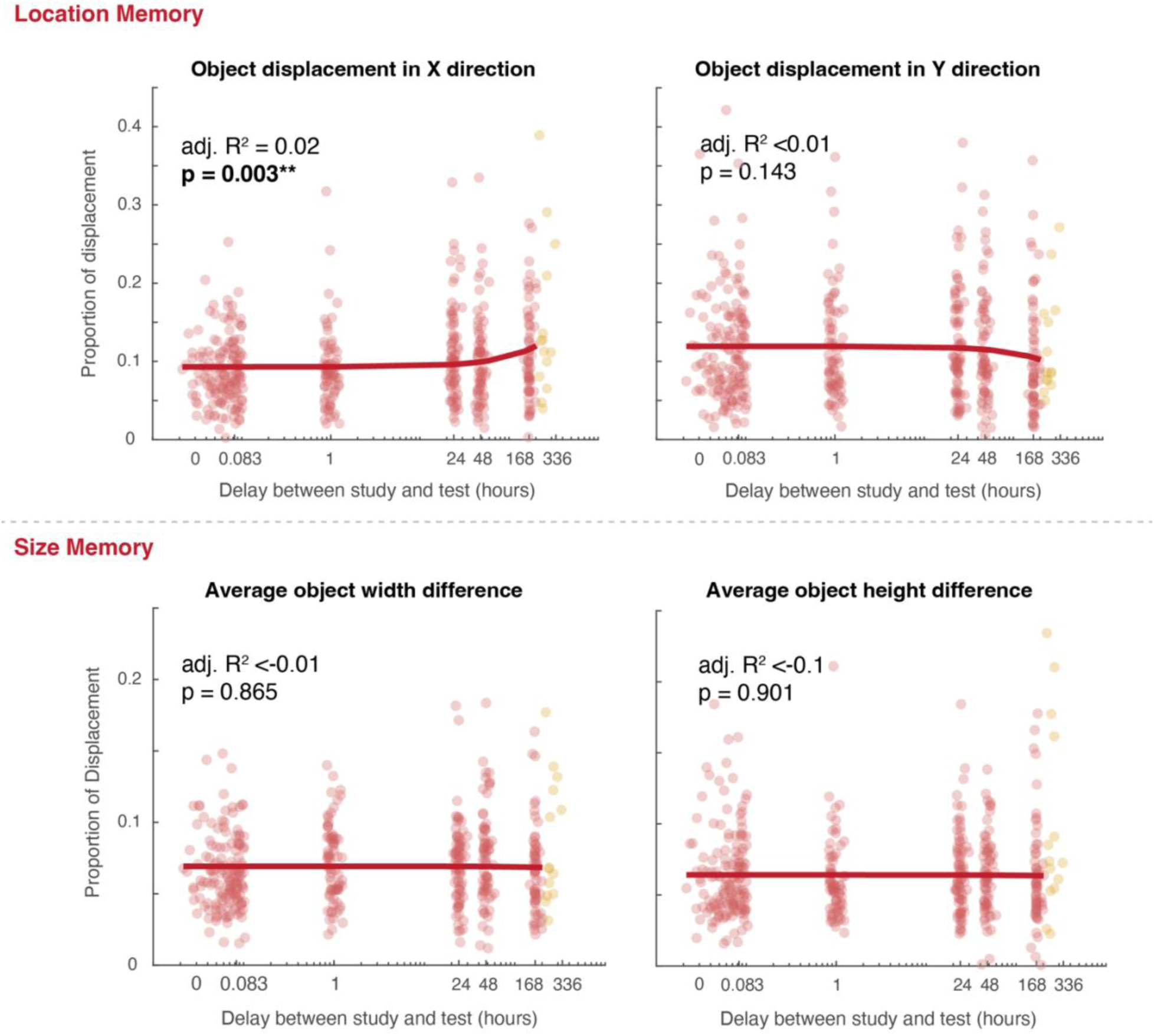
Linear regressions predicting spatial accuracy from delay time. All x-axes are in the logarithmic scale. Each red dot is a single drawing, and each yellow dot is a drawing from the group of participants that returned 1-2 weeks after encoding. Although these yellow dots are visually included, they were not included in the analyses. (top panel) For location memory, delay time was only significantly predictive of location accuracy in the x-direction, but not the y-direction, meaning that increased delay time led to worse location memory in the x-direction. (bottom panel) Unlike location memory, size memory was unaffected by delay time, with neither width memory nor height memory influenced by how much time elapsed before recall.

We also determined the features of these false objects by identifying trends in their identities as well as their locations within the drawings. False objects tended to be objects semantically related to the scene recalled, such as a swingset (5 drawings) or merry-go round in the playground (1 drawing), a clock in the living room (2 drawings), or flowers on the kitchen table (2 drawings). Interestingly, there were also many false weather conditions (10 drawings), like the sun (4 drawings) or clouds (4 drawings), despite the absence of the sky in most of the images. When we covered the false objects with ellipses to create a heatmap of their locations (see **Fig. 9c**), we found that though false objects occasionally shared the location of true objects from the recalled image, there were overall few observable trends for the location of false objects. Overall, these results are consistent with the idea of our memories becoming more semantic (or schematic) over delay time, resulting in an increase in false objects that are related to the scene being recalled.

### Spatial memory precise even one week later

In additional to *what* is in memory, is our memory for *where* also affected by how much time has elapsed between encoding and recall? To test this, we had 5 AMT workers (831 workers total) determine the size (width and height) and location (*x*-direction location and *y*-direction location) of every recalled object by moving and resizing an ellipse to cover each object.

Participants overall performed well when recalling the location of objects from a scene, displacing objects on average by 9.78% (*SD* = 5.30%) of the image in the *x*-direction and by 11.62% (*SD* = 6.94%) of the image in the *y*-direction. We found that *delay time* did significantly positively predict (p = 0.003) the *displacement of objects in the x-direction* using a linear regression (F(426) = 9.09, p = 0.003, adjusted *R*^2^ = 0.02), meaning that participants worsened in their location memory as more time elapsed between encoding and recall. However, we did not find this same effect in the *y*-direction; if anything, location memory in the *y*-direction actually improved over delay time, although this was not significant (F(426) = 2.15, p = 0.143, adjusted *R*^2^ <0.01).

When recalling the size of objects from the encoded scene, participants were extremely accurate, and were only off by 6.91% (*SD* = 2.85%) of the image for object width and 6.40% (*SD* = 3.04%) of the image for object height. Unlike location memory, delay time had no effect for size memory. Using separate linear regressions, we found that delay time did not significantly predict the accuracy of width memory (F(426) = 0.029, p = 0.865, adjusted *R^2^* <-0.01) or height memory (F(426) = 0.016, p = 0.901, adjusted *R^2^* <-0.01) when tested using two separate linear regressions. Indeed, even one-week after encoding, participants on average were only off by 6.65% (*SD* = 3.06%) of the image for object width and 6.33% (*SD* = 3.62%) of the image for object height.

Although these results indicate that delay time appears to have little effect on the accuracy of spatial memory, it is possible that other factors can explain some of the variance in accuracy between participants. Therefore, we tested whether spatial imagery ability—determined through the Spatial score of the OSIQ—correlates with the accuracy of location memory. We chose location memory as it is more classically viewed as spatial memory than size memory. We measured location error as the Euclidean distance between the centroid of the drawn object and the centroid of the object in the photograph in both the *x*- and *y*-directions.

We first correlated participants’ OSIQ-Spatial scores to their spatial memory across all time conditions (see **Fig. 11** left panel), but this correlation was not significant (Spearman’s rank correlation: ρ <0.01, p = 0.950). As time significantly impacted the accuracy of location memory in the *x*-direction, we ran a second correlation limited to the three shortest time conditions in which the change in time was minimal (no delay – 1 hour delay; see **Fig. 11** right panel), but there was similarly no correlation between spatial imagery ability and location memory (Spearman’s rank correlation: ρ = −0.05, p = 0.506). These results suggest that participants overall were surprisingly accurate in both their location and size memory across delays, even having highly accurate spatial memory one-week after encoding. But that, unlike object imagery, spatial imagery ability does not appear to influence the accuracy of spatial memory.

**Figure 11.**
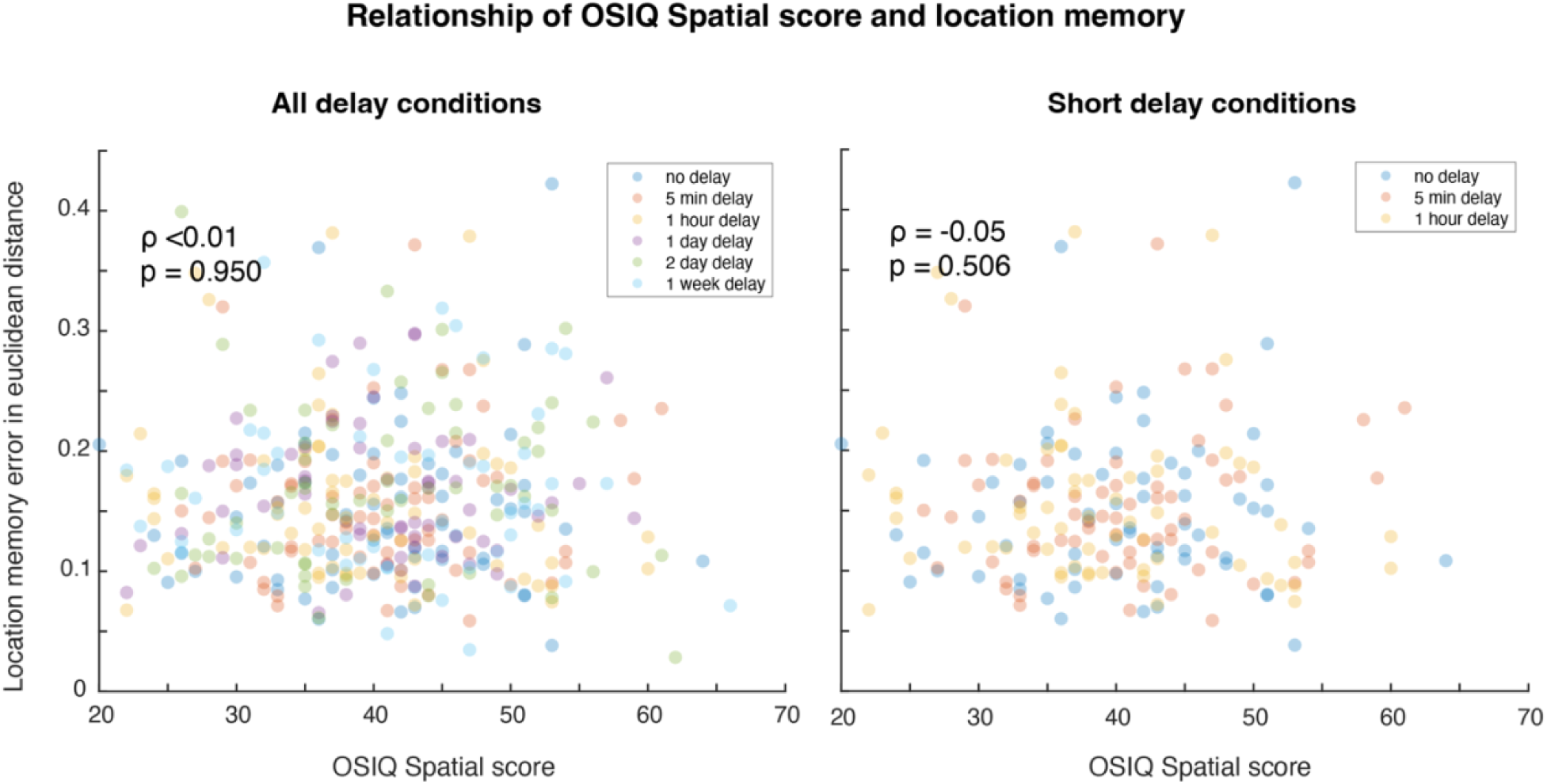
Relationship of spatial imagery ability and location memory. (left panel) Correlation between location memory, measured as error in Euclidean distance, and the Spatial score on the OSIQ, across all time delays. We found no significant correlation between spatial imagery and spatial memory. (right panel) Correlation between location memory and the OSIQ Spatial score for the three shortest delay times, in which there were relatively minimal changes in time. There was similarly no significant correlation between spatial imagery and spatial memory.

In total, we found that participants overall had surprisingly accurate memory for a scene even one-week after encoding. In fact, spatial memory was largely unaffected by delay time, with only memory for the location of objects in the *x*-direction worsening across delay. In contrast, we found that object memory was impacted by time, with participants recalling a significantly lower proportion of objects from a scene with longer periods of delay between encoding and recall, as well as more false objects. We also found that the saliency of an object, rather than its location or meaning, predicted when an object would be forgotten, with objects low in saliency the first forgotten. Lastly, we found that whereas object imagery ability interestingly influences the proportion of objects recalled from a scene, spatial imagery ability did not appear to influence spatial memory in the same way.

## General Discussion

In this work, we show that hundreds of drawings created from memory across ranges of encoding durations (Experiment 1) and retention periods of the memory (Experiment 2) were able to holistically capture the features of memory that are built up and broken down across time. Online crowdsourced scoring of these drawings revealed that the proportion of objects recalled from a scene and the presence of falsely recalled objects not in the original scene were highly dependent on both encoding and delay time. In contrast to the objects recalled, we found that spatial memory was largely unaffected by time, with surprisingly accurate memory for the location and size of objects after minimal encoding of 100 msec or a 1-week delay between encoding and recall. Additionally, we found that different object features predicted which objects were recalled across time, with meaningful and central objects having an advantage of being encoded into memory first, and objects low in visual saliency being the first objects forgotten. Lastly, we found that whereas higher object imagery ability resulted in a higher proportion of objects recalled, spatial imagery ability did not affect spatial memory in the same way.

These results were able to replicate large, classic findings investigating the effect of time on memory despite our use of different methods and stimuli. We were even able to parallel Ebbinghaus’ classically dubbed forgetting curve (1885), in which he found a decrease in the words remembered from a word list as time increased before recall. Here, we found similar patterns of forgetting within a *single visual stimulus*, with participants drawing fewer objects from a scene as more time passed since encoding. Additionally, we replicated prior studies utilizing recognition, as opposed to recall, finding more objects remembered from a scene across encoding (Melcher, 2001, 2006), as well as changes in the reliability of memory (i.e., likelihood of false memories) across time (Lampinen et al., 2001; Memon et al., 2003). In addition to verifying that these classical findings hold across different measurements of memory, these replications also provide evidence that using drawing recall can accurately capture the content of memory.

Beyond replicating classical findings, the detail contained within drawings was also able to quantify memory across time in new and exciting ways. Although there have been muddied results on how spatial memory changes over the course of encoding (Melcher, 2006; Tatler et al., 2003) and forgetting (Mandler & Ritchey, 1977; Talamini & Gorree, 2012) in recognition studies, our results provide evidence that memory for both the location and size of objects remains accurate and relatively unchanged across these processes. As this directly contrasts with our findings that object memory is highly dependent on time, this suggests that there are differing effects of time on these memory components. Indeed, our findings contribute to a growing body of work suggesting a dissociation between spatial and object memory. An interesting case study provides evidence that one memory component can be impaired with the other intact, with a patient MV having impaired localization of objects in working memory, but preserved working memory for their visual features (Carlesimo et al., 2001). Likewise, those with *aphantasia*, a condition characterized by the lack of voluntary mental imagery (Keogh & Pearson, 2018; Zeman et al., 2015), have been found to have intact spatial memory despite disrupted object memory (Bainbridge, Pounder, et al., 2021). A dissociation between spatial and object memory may be explained by the involvement of different brain regions in these memories, as the parahippocampal cortex may play a role in spatial memory, but the perirhinal cortex in object memory (Diana et al., 2007, 2012; Reagh & Yassa, 2014; Staresina et al., 2011).

The present results do not only provide further evidence for a dissociation between spatial and object memory, but also within spatial memory itself. Across encoding time, we found that whereas memory related to the *x*-coordinates of objects (i.e., location of objects in the *x*-direction and the width of objects) did not significantly change, memory related to the *y*- coordinates of objects did. Similarly, we found that whereas participants worsened in their location memory of objects in the *x*-direction across delay time, participants’ location memory in the *y*-direction remained unchanged—or if anything, improved—across delays in recall.

In other words, we observed differences in spatial memory for *x*- and *y*-coordinates across both encoding and delay time. Although we believe these are novel results in the time and memory literature, brain regions involved in scene processing, such the parahippocampal place area (PPA) and transverse occipital sulcus (TOS), have been reported to have differential activation biases for the upper and lower visual fields (Silson et al., 2015). Therefore, it would be valuable for future work to further evaluate whether these spatial biases during scene processing are reflected in how memory changes across time.

Collectively, our results are consistent with both gist-based processing over encoding (Ahmad et al., 2017a; M. R. Greene & Oliva, 2009; Oliva, 2005) and the generalization of memory over delay (Harvey, 1986; Posner & Keele, 1970; Sweegers & Talamini, 2014; Zeng et al., 2021). We find that when participants only had a brief time to encode an image, they were likely only able to extract the ‘gist’ or semantic label of an image, resulting in the recall of objects high in meaning, scene-consistent false objects, but overall few details from the scene. Future work using eye-tracking during encoding could offer valuable insight into whether the accumulation of image-specific details correlates with participants’ ability to explore the scene. Similarly, we find evidence for the generalization of memory over delay time, as participants seemingly revert back to a more semantic label of the image as more time passes before recall. This is supported by the findings of participants first forgetting objects due to their saliency, rather than their meaning, recalling more false objects semantically consistent with the recalled scene over time, and participants forgetting more objects over time. Interestingly, the parallel found in the current study between gist-based processing and the generalization of memory supports the idea that the first information in memory may also the last to leave (Potter, 2012), despite other recent studies challenging this idea (Goujon et al., 2022).

The differences we found in memory across time are applicable to real-world scenarios such as eyewitness testimony, with our results suggesting that some features of memory, such as spatial memory, may be more accurate during testimony. However, it is possible that we found such accurate and consistent memory across encoding and delay time because we used organized scenes, in which participants could potentially utilize their pre-existing knowledge to guide the size and location of objects. Indeed, previous studies have found better spatial memory for organized than disorganized scenes (Draschkow & Võ, 2017), and even that spatial memory worsens more quickly over delay time if scenes are not arranged in a meaningful way (Mandler & Parker, 1976; Mandler & Ritchey, 1977). Additionally, studies have found that spatial memory may take the largest dip in accuracy after longer delays than those used in the present study, such as one to three months after encoding (Talamini & Gorree, 2012). This is an important consideration as these longer delay times are more akin to when eyewitness testimony would take place (Schacter & Loftus, 2013). Therefore, future work should test the applicability of our findings in ways that may better transfer to these more real-world scenarios.

In conclusion, the rich content captured in drawings made from memory was able to holistically and uniquely characterize the features of memory that change with time. Comparing the large, extending effect of time on object memory to the minimal (or no) effect of time on spatial memory suggests a dissociation in our memory for what and where, even within a single scene image.

## Data and code availability

All data, including participant drawings, will be available upon publication on Open Science Framework (OSF).

## Acknowledgements

We would like to thank Rebecca Greenberg for running the initial encoding time drawing experiment as well as Asaf Lebovic for his assistance scoring a portion of these drawings. The present work was supported by the National Eye Institute (R01-EY034432) to W.A.B.

