## Supplementary Materials for "Drawings reveal changes in object memory, but not spatial memory, across time"

301 Green Hall

5848 S. University Ave.

Chicago, IL 60637

Contents Page

**Supplemental Table 1.1:** Model statistics for the seven Experiment 1 linear regression models 3

**Supplemental Table 1.2:** Model statistics for the Experiment 1 logistic regression model 5

**Supplemental Table 1.3:** Model statistics for the seven Experiment 2 linear regression models 6

**Supplemental Table 1.4:** Model statistics for the Experiment 2 multiple linear regression 8

**Supplemental Table 1.5:** Model statistics for the Experiment 2 logistic regression model 9

**Supplemental Table 1.1**

*Model statistics for the seven Experiment 1 linear regression models*

| Model, designated by dependent variable | *Encoding Time β estimate* | *Adjusted $R^2$* | *p-value* | *Akaike Information Criterion (AIC)* |
|---|---|---|---|---|
| Proportion of objects recalled | $1.05 \times 10^{-5}$ | 0.34 | <0.001*** | -796 |
| Number of false objects | $-1.10 \times 10^{-5}$ | 0.13 | 0.027* | 270 |
| Displacement of objects in *x*-direction location | $3.05 \times 10^{-8}$ | 0.02 | 0.980 | -721 |
| Displacement of objects in *y*-direction location | $-2.30 \times 10^{-6}$ | 0.04 | 0.035* | -791 |
| Displacement of object *width* | $-9.26 \times 10^{-7}$ | 0.06 | 0.206 | -1053 |
| Displacement of object *height* | $-1.43 \times 10^{-6}$ | 0.11 | 0.037* | -1049 |

*Note.* β values are low because of the scale difference between the dependent (range: 0 – 1) and independent variables (range: 100 msec – 10000 msec).

*Formulas for the seven Experiment 1 linear regression models*

**Proportion of objects recalled:**

$$proportion\ of\ objects\ recall \sim \beta_0 + \beta_1(encoding\ time) + (encoding\ time \mid participant)$$

**Number of false objects:**

$$number\ of\ false\ objects \sim \beta_0 + \beta_1(encoding\ time) + (encoding\ time \mid participant)$$

**Displacement of objects in *x*-direction:**

$$displacement\ of\ objects\ in\ x\text{-}direction \sim \beta_0 + \beta_1(encoding\ time) + (encoding\ time \mid participant)$$

**Displacement of objects in *y*-direction:**

$$displacement\ of\ objects\ in\ y\text{-}direction \sim \beta_0 + \beta_1(encoding\ time) + (encoding\ time \mid participant)$$

**Displacement of object *width*:**

$$displacement\ of\ object\ width \sim \beta_0 + \beta_1(encoding\ time) + (encoding\ time \mid participant)$$

**Displacement of object *height*:**

$$displacement\ of\ object\ height \sim \beta_0 + \beta_1(encoding\ time) + (encoding\ time \mid participant)$$

**Supplemental Table 1.2**

*Model statistics for the Experiment 1 logistic regression model*

| Statistic | *Encoding Time* | *Object Meaning* | *Object Saliency* | *Encoding Time : Object Meaning* | *Encoding Time : Object Saliency* |
|---|---|---|---|---|---|
| β Estimate | 0.37 | 0.38 | 0.52 | -0.14 | -0.01 |
| *p-value* | <0.001*** | <0.001*** | <0.001*** | 0.008** | 0.781 |
| | | | *Adjusted $R^2$* | *p-value* | *AIC* |
| Overall Model Statistics: | | | 0.05 | <0.001*** | 51578 |

*Note*. AIC stands for Akaike information criteria.

**Model formula:**

$$\begin{aligned} object\ recalled \sim \beta_0 + \beta_1(encoding\ time) + \beta_2(object\ meaning) + \beta_3(object\ saliency) \\ + \beta_4(encoding\ time{:}\ object\ meaning) \\ + \beta_5(encoding\ time{:}\ object\ saliency) + (1 \mid Drawing) \end{aligned}$$

* Note that because the meaning and saliency values were correlated, we residualized these values before using them in the model.

**Supplemental Table 1.3**

*Model statistics for the seven Experiment 2 linear regression models*

| Model, designated by dependent variable | *Delay Time* β *estimate* | *Adjusted* $R^2$ | *p-value* | *Akaike Information Criterion (AIC)* |
|---|---|---|---|---|
| Proportion of objects recalled | -0.0006 | 0.06 | <0.001*** | -418 |
| Number of false objects | 0.0012 | 0.02 | <0.001*** | 624 |
| Displacement of objects in *x*-direction location | 0.0001 | 0.02 | 0.003** | -1306 |
| Displacement of objects in *y*-direction location | $-8.57 \times 10^{-5}$ | <0.01 | 0.143 | -1068 |
| Displacement of object *width* | $4.10 \times 10^{-6}$ | <-0.01 | 0.865 | -1829 |
| Displacement of object *height* | $-3.20 \times 10^{-6}$ | <-0.01 | 0.901 | -1772 |

*Note.* β values are low because of the scale difference between the dependent (range: 0 – 1) and independent variables (range: 0 hr –168 hr).
*Note II*. Formulas for these seven linear regression models are shown below.

**Proportion of objects recalled:**

$$proportion\ of\ objects\ recall \sim \beta_0 + \ \beta_1(delay\ time)$$

**Number of false objects:**

$$number\ of\ false\ objects \sim \beta_0 + \ \beta_1(delay\ time)$$

**Displacement of objects in *x*-direction:**

$$displacement\ of\ objects\ in\ x\text{-}direction \sim \beta_0 + \ \beta_1(delay\ time)$$

**Displacement of objects in *y*-direction:**

$$displacement\ of\ objects\ in\ y\text{-}direction \sim \beta_0 + \ \beta_1(delay\ time)$$

**Displacement of object *width*:**

$$displacement\ of\ object\ width \sim \beta_0 + \ \beta_1(delay\ time)$$

**Displacement of object *height*:**

$$displacement\ of\ object\ height \sim \beta_0 + \ \beta_1(delay\ time)$$

**Supplemental Table 1.4**

*Model statistics for the Experiment 2 multiple linear regression*

| Statistic | *Base model* | *Interaction model* |
|---|---|---|
| *Delay Time* β *Estimate* | -0.29*** | -0.29*** |
| *Object Score* β *Estimate* | 0.10* | 0.10* |
| *Delay Time : Object Score* β *Estimate* | — | 0.02 |
| *Adjusted* $R^2$ | 0.09 | 0.09 |
| *p-value* | <0.001*** | <0.001*** |
| *Akaike Information Criterion (AIC)* | 1058 | 1060 |

*$p<0.05$. **$p<0.01$. ***$p<0.001$.

**Base model formula:**

$$proportion\ of\ objects\ recall \sim \beta_0 + \beta_1(delay\ time) + \beta_2(objectScore)$$

**Interaction model formula:**

$$proportion\ of\ objects\ recall \sim \beta_0 + \beta_1(delay\ time) + \beta_2(objectScore) + \beta_3(delay\ time : objectScore)$$

**Supplemental Table 1.5**

*Model statistics for the Experiment 2 logistic regression model*

| *Statistic* | *Delay Time* | *Object Meaning* | *Object Saliency* | *Delay Time : Object Meaning* | *Delay Time : Object Saliency* |
|---|---|---|---|---|---|
| β Estimate | -0.21 | 0.09 | 0.28 | -0.03 | -0.09 |
| *p-value* | <0.001*** | 0.0163* | <0.001*** | 0.484 | 0.023* |
| | | | *Adjusted $R^2$* | *p-value* | *AIC* |
| Overall Model Statistics: | | | 0.02 | <0.001*** | 47376 |

*Note*. AIC stands for Akaike information criteria.

**Model formula:**

$$object\ recalled \sim \beta_0 + \beta_1(delay\ time) + \beta_2(object\ meaning) + \beta_3(object\ saliency) + \beta_4(delay\ time\colon object\ meaning) + \beta_5(delay\ time\colon object\ saliency)$$

* Note that because the meaning and saliency values were correlated, we residualized these values before using them in the model.
